# Tracing Autism Traits in Large Multiplex Families to Identify Endophenotypes of the Broader Autism Phenotype

**DOI:** 10.1101/659722

**Authors:** Krysta J Trevis, Natasha J Brown, Cherie Green, Paul Lockhart, Peter Hickey, Miriam Fanjul-Fernández, Catherine Bromhead, Tarishi Desai, Tanya Vick, Greta Gillies, Hayley Mountford, Elizabeth Fitzpatrick, Lavinia Gordon, Peter Hewson, Vicki Anderson, Martin B Delatycki, Ingrid E Scheffer, Sarah J Wilson

## Abstract

Families comprising many individuals with Autism Spectrum Disorder (ASD) may carry a dominant predisposing mutation. Our aim was to use rigorous phenotyping of the ‘Broader Autism Phenotype’ (BAP) in large multiplex ASD families to identify endophenotypes of the BAP for future genetic studies. We evaluated ASD/BAP features using standardised tests and a semi-structured interview to assess social, intellectual, executive and adaptive functioning in 109 individuals, including two large multiplex families (Family A: 30; Family B: 34) and an independent sample of small families (*n*=45). Our protocol identified four psychological endophenotypes of the BAP that were evident in both samples, and showed high sensitivity (97%) and specificity (82%) for individuals classified with the BAP. The patterns of inheritance of these endophenotypes varied in the two large families, supporting their utility for identifying genes in autism.

---

Autism Spectrum Disorder (ASD) is a neurodevelopmental condition that spans deficits in two domains: social communication, and restricted interests or repetitive behaviours APA 2013. Recent estimates from the Center for Disease Control and Prevention indicate prevalence of ASD is one in 68 children aged 8 years^1^ making ASD a critical international health problem. Clinical and molecular research provides evidence for a genetic aetiology in ASD, yet despite recent molecular advances the cause remains unidentified in the majority of cases.

Improved understanding of ASD has facilitated recognition of milder phenotypes. Early clinical research identified autistic traits in relatives of children with ASD, described as the ‘broader autism phenotype’ (BAP)^2,3^. The BAP sits at the mildest end of the ASD spectrum and includes a range of subtle behavioural and cognitive features that reflect the two core domains of ASD. The *Diagnostic and Statistical Manual of Psychiatric Disorders* –*V* (DSM-V) stipulates that a significant degree of impairment must be present to qualify for a diagnosis of ASD^4^, whereas BAP traits lie on a continuum of normal population behaviours^5^. Monozygotic twins demonstrate 30% concordance for ASD^6^, increasing to 92% if the BAP is considered, while dizygotic twin concordance is ∼10%^7^. Prevalence of the BAP in the general population is unknown, whereas several studies have demonstrated higher rates of BAP traits (20-50%) among relatives of children with ASD compared to controls, particularly in the areas of pragmatic language^8,9^, personality^10^, social cognition^9,11,12^ and executive function^13,14^. Together these findings support the notion of complex inheritance of ASD.

The overall diagnostic rate of ASD is now >30%, including monogenic and chromosomal aetiologies^15,16^. The remaining ∼70% are likely to have a genetic basis, with polygenic architecture in some, and unique *de novo* mutations in others^17^. Complementary techniques will be necessary to unravel aetiology in unsolved cases. Typically, family studies combine many small families (2-3 affected individuals), however, these are likely to be confounded by genetic heterogeneity. Very large multiplex families (> 8 affected) where ASD traits appear dominantly inherited are rare, but more genetically homogeneous. In other complex disorders, such as epilepsy, phenotypic characterisation of such families has proved powerful in gene discovery^18^, however this approach has received limited attention in ASD^1818^. In multiplex ASD families, the identification of family members with BAP traits, or endophenotypes, may serve as markers of carrier status^19,20^. In turn, this may facilitate gene identification^21^.

Endophenotypes are measurable features within a disorder that are proposed to reduce its complexity into more quantifiable elements^22^. They have been hypothesised to reflect more aetiologically homogeneous subgroups within genetically heterogeneous conditions. There are several BAP traits that may be considered “endophenotypes” from within the domains of language, executive function, and social cognition^21^. In the context of a single large family where numerous individuals demonstrate ASD or the BAP, recognition of BAP endophenotypes should allow granular identification of an autism gene of dominant effect. This study is the first known to the authors to apply this approach in autism.

The aim of our study was to analyse autistic traits within large multiplex families to examine inheritance of ASD by identifying endophenotypes of the BAP. We achieved this aim using an iterative process, first rigorously phenotyping many members of large multiplex families to delineate the full range of BAP traits for potential endophenotypes. We then assessed these traits in a separate sample of 20 small families, each with at least one member with ASD, to independently validate the endophenotypes. We then applied these endophenotypes to two of our fully characterised large multiplex families (from step 1) to assess their utility for examining inheritance patterns. We hypothesised that (1) multiple individuals in large families would demonstrate the BAP, (2) specific BAP endophenotypes would be identifiable across the traditional BAP domains, and (3) these endophenotypes would vary in presentation between large multiplex families.

## Methods

### Large Multiplex Families

Large multiplex families were primarily ascertained from the Barwon Autism Database as part of a broader Collaborative Autism Study^23^. For inclusion as a multiplex family, > 8 individuals with a diagnosis or suspected diagnosis of ASD or the BAP were required. The two fully characterised large multiplex families used to examine inheritance patterns using BAP endophenotypes are referred to as ‘Family A’, ascertained from the Barwon Autism Database, and ‘Family B’, who was self-referred. All available relatives were recruited, including those with and without reported BAP traits. Informed consent was obtained from all participants or a parent/guardian, following approval of the study by the Human Research Ethics Committees of Barwon Health and the Royal Children’s Hospital, Melbourne.

#### Protocol for Diagnosing ASD in Large Multiplex Families

ASD diagnoses were confirmed using the Autism Diagnostic Observation Schedule-Generic^24^ (ADOS-G), the Autism Diagnostic Interview-Revised^25^ (ADI-R), or DSM-IV-TR criteria^26^. For adults, the structured Family History Interview^2^ (FHI) was administered by NJB, while for adolescents, a detailed developmental and medical history was obtained. Quantitative measures of intellect, executive functions, adaptive behaviour and social functioning were also completed (Table 1). Testing was undertaken over a number of days to minimise fatigue effects. A physical examination was conducted for dysmorphic and neurocutaneous features and growth parameters. Standard genetic testing (karyotype, fragile X testing) and metabolic investigations were performed on probands.

**Table 1.** Protocol for diagnosing ASD and phenotyping the BAP in large multiplex families.

| Protocol Item | Participants with ASD |  | Participants without ASD |  |  |
| --- | --- | --- | --- | --- | --- |
|  | Child or Adolescent<br>≥4.5-17yr | Adult<br>≥18yr | Child<br><13yr | Adolescent<br>≥13–17yr | Adult<br>≥18yr |
| ADI-R + ADOS-G *or* DSM-IV interview + ADOS-G | + | ± | - | - | - |
| Detailed developmental, medical, psychiatric and behavioural history | + | + | + | + | + |
| Family History Interview | - | + | - | - | ± |
| Standardised testing of cognition and executive function<sup>a</sup> | ± | + | + | + | ± |
| Questionnaires of adaptive behaviour<sup>b</sup> | + | + | + | + | ± |
| Broader Autism Phenotype Interview, the Faux Pas Task, Cartoon Task and Pragmatic Rating Scale | - | ± | - | + | + |
| Physical Examination | + | + | + | + | + |
| High resolution molecular karyotype, Fragile X testing, metabolic investigations | + | - | - | - | - |

*ASD* Autism Spectrum Disorder, *ADI-R* Autism Diagnostic Interview-Revised, *ADOS-G* Autism Diagnostic Observation Schedule-Generic, *DSM-IV* Diagnostic and Statistical Manual of Mental Disorders (4th edition).
+ all individuals completed this assessment; - no individuals completed this assessment; ± only some individuals completed this assessment
<sup>a</sup>Wechsler Abbreviated Scale of Intelligence and subtests of the Delis-Kaplan Executive Function System
<sup>b</sup>The Adaptive Behavioural Assessment System (2nd edition) and The Behavioural Rating Inventory of Executive Function

Across Family A and B, 65 individuals were recruited: 16 children (2-12 years), 9 adolescents (13-17 years) and 40 adults (18-79 years) spanning 4 generations. Of these, 16/65 met criteria for a diagnosis of ASD. Family B also reported a deceased family member who had a diagnosis of ASD (generation V) and an additional family member with ASD (generation 4) who was not recruited. Scrambled pedigrees of affected status are presented to preserve participant anonymity (Fig.1). In each family, a matriarch was identified. Individuals directly related to each matriarch are classified as ‘core family’; others are referred to as ‘married-in’. Family A comprised 30 individuals, including 7 diagnosed with ASD (6/9 children, 1/6 adults; Fig.1a). Nineteen were core family; 11 were married-in. In Family B, we fully phenotyped 35 individuals, not including the matriarch (who was not assessed). Three children and three adolescents participated in a limited range of phenotyping activities and as such, these individuals were excluded from final analyses. Nine had ASD (5/7 children, 1/5 adolescents, 3/22 adults; Fig.1b); 31 were core family, and four were married-in.

**Figure 1.**
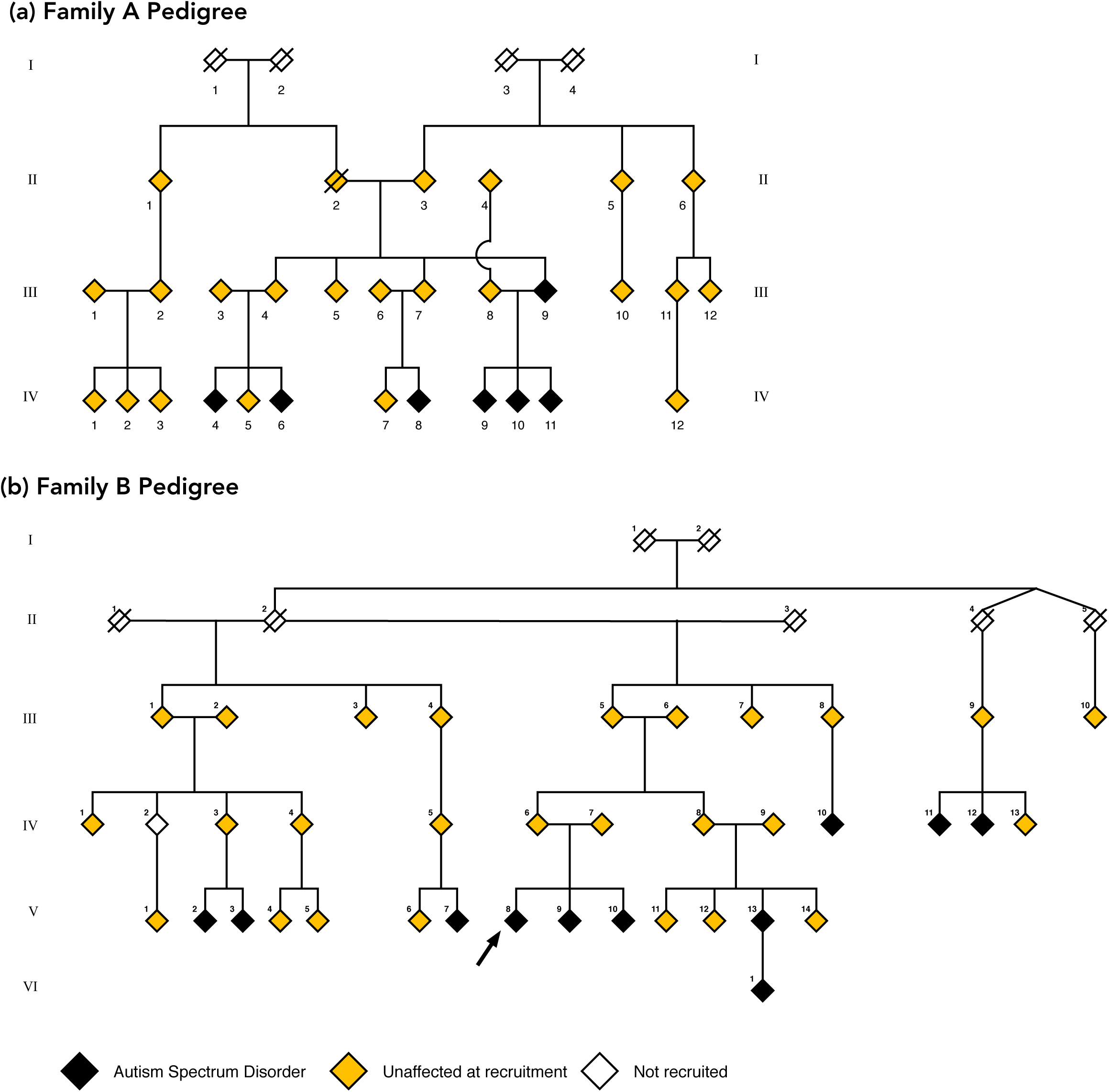
Scrambled pedigrees for Family A (panel A) and Family B (panel B) at recruitment. Individuals with a diagnosis of ASD are marked in black, and individuals recruited from the broader families are marked in yellow. White diamonds are individuals who were not recruited but are represented here to preserve the pedigree lines.

#### Protocol for Phenotyping the BAP in Large Multiplex Families

We employed a mixed methods approach to rigorously assess the BAP, including an evaluation of general intellect, executive functions, adaptive behaviour, social cognition and language pragmatics (Table 1). A purpose developed semi-structured interview, the Broader Autism Phenotype Interview (BAPI), was also administered by three clinicians with expertise in neurobehavioural disorders (NJB, SJW, IES) to all individuals ≥13 years, to determine the presence, nature and extent of BAP features. Questions focused on the participant’s life story, personal qualities, relationships, social functioning, and developmental, medical, psychiatric and vocational history. During the interview we included one or two “intentional errors” to elicit pragmatic elements of the BAP, such as terse speech^9^.

Full scale (FSIQ), verbal (VIQ) and performance (PIQ) intelligence quotients were derived with the four subtest Weschler Abbreviated Scale of Intelligence^27^ (WASI; *M*=100, *SD*=15). Executive functions were measured with seven subtests of the Delis-Kaplan Executive Function System^28^ (D-KEFS; *M*=10, *SD*=3). The second edition of the Adaptive Behavioural Assessment System^29^ (ABAS-II) and the Behavioural Rating Inventory of Executive Function^30^ (BRIEF) were used to assess adaptive functioning (Table 1).

Social discourse was assessed using an adapted Faux Pas Task^31,32^ (FPT). The task included four faux pas stories and four control stories^33^ (maximum score=40, *M*=37, *SD*=4). In addition, the Goldman-Eisler Cartoon task^34^ was used to explore previous observations of overly detailed speech and longer pauses between words in the BAP^9^. This task measures discourse production by eliciting a description of an eight frame captionless cartoon, “The Cowboy Story”, over three successive trials^35^. Control individuals show increased verbal fluency with successive trials compared with decreased fluency in individuals with communication deficits^34^. Following all assessments, the Pragmatic Rating Scale (PRS) was independently completed by the three interviewers and consensus ratings reached. A score ≥4 defined pragmatic impairment^10^. After independent review of all qualitative and quantitative data by NJB, SJW, and IES the presence of the BAP was determined by consensus.

### Small Families

We recruited an independent sample of 45 individuals from 20 small families with at least one member diagnosed with ASD, via advertisements and from the Barwon Autism Database. All participants provided written informed consent, as described above. Inclusion criteria were: (i) no diagnosis of ASD (based on DSM-IV or DSM-V criteria), (ii) ≥1 family member with ASD (based on DSM-IV or DSM-V criteria), and (iii) >12 years of age. Individuals were classified as having the BAP if they met ≥2 criteria for a BAP diagnosis on the Broader Autism Phenotype Rating Scale^2^ (BAPRS). Individuals were classified as unaffected if they did not meet criteria for any BAP traits or a diagnosis of ASD. This identified 30 individuals with the BAP (4 adolescents, 26 adults) in the 20 families, ranging in age from 14-71 years, and 11 unaffected adult family members ranging in age from 18-53 years. Four adult individuals showed only one BAP trait on the BAPRS and thus, were excluded from analyses based on the above criteria.

In 27 individuals, average total scores were available for the Broader Autism Phenotype Questionnaire (BAPQ), and in 31 individuals, FSIQ, Verbal Comprehension (VCI) and Perceptual Reasoning (PRI) indices had been derived with the WASI-II^36^ (*M*=100, *SD*=15). As shown in Table 2, all individuals were within the normal range based on FSIQ, with no significant differences between unaffected and BAP individuals for age or intellect (all *p*<.250). Consistent with expectations, there was a trend for higher scores on the BAPQ in the BAP group, with a medium effect size (*t*(24.56)=-1.96, *p*=.062, *d*=0,70).

**Table 2.** Demographics of the small families sample.

|  | Unaffected | BAP |
| --- | --- | --- |
| Number of participants (female) | 11 (6) | 30 (19) |
| Mean age (range) | 41.09 (18-53) | 39.50 (14-53) |
| Mean BAPQ (SD)* | 2.43 (0.38) | 2.92 (0.92) |
| Mean FSIQ (SD)** | 108 (14) | 111 (13) |
| Mean VCI (SD)** | 106 (18) | 109 (15) |
| Mean PCI (SD)** | 109 (7) | 110 (15) |

\*data available for unaffected ( $n=9$ ) and BAP ( $n=18$ ); \*\* data available for unaffected ( $n=10$ ) and BAP ( $n=21$ ). BAPQ=Broader Autism Phenotype Questionnaire; FSIQ=Full Scale Intelligence Quotient; VCI=Verbal Comprehension Index; PRI=Perceptual Reasoning Index

### Procedure

We used an iterative process to characterise, refine and assess endophenotypes of the BAP in our two separate samples, as summarised in Fig.2.

**Figure 2.**
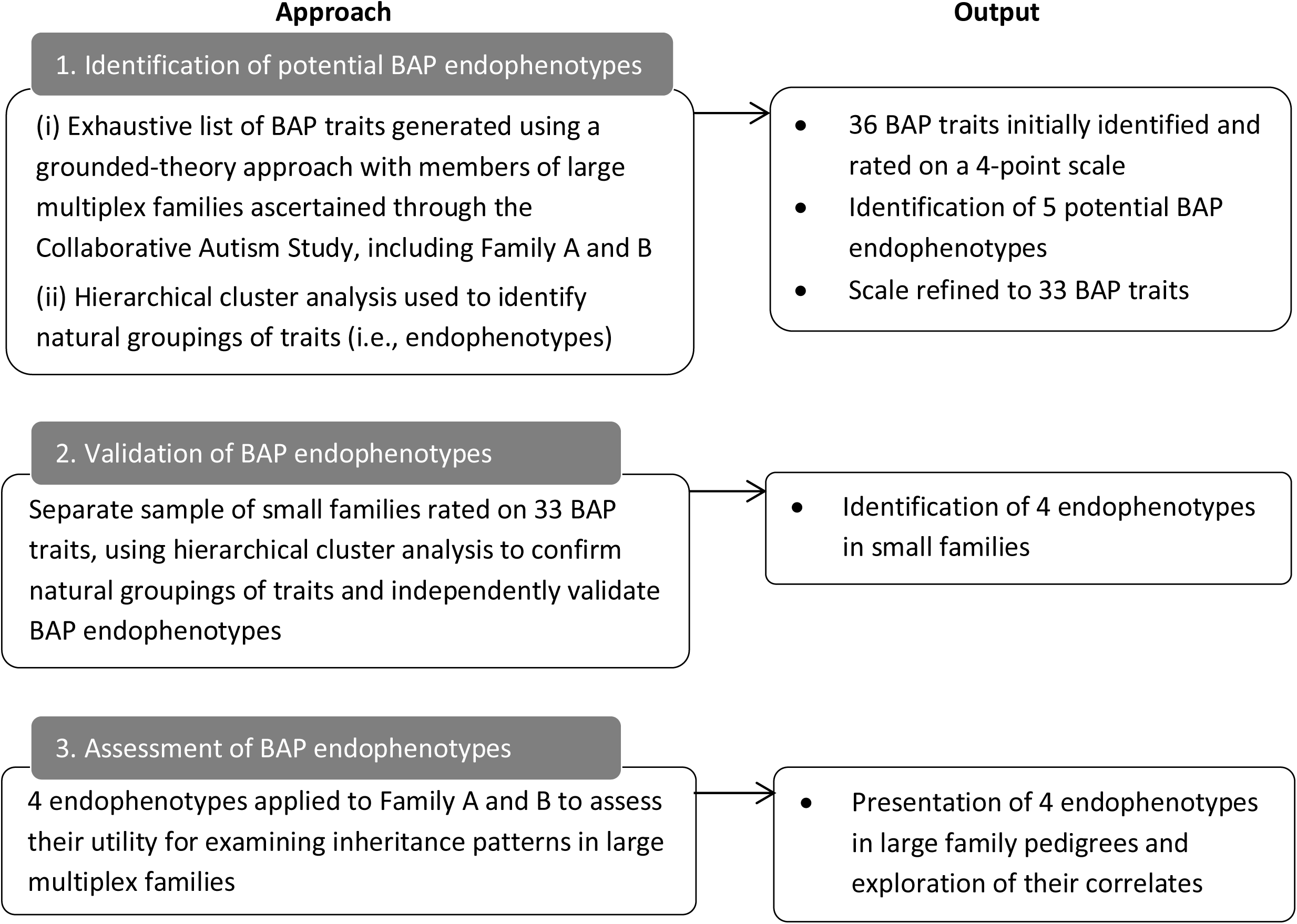
Iterative process used to identify and assess BAP endophenotypes.

#### Step 1: Identification of Potential BAP Endophenotypes in Large Multiplex Families

Using a grounded theory approach, BAP traits were initially identified from a detailed literature review targeting the theoretical domains described in the seminal work of Bolton (1994), on which the conceptualisation of the BAP is largely based. The domains included speech, literacy, pragmatics, relationships, and circumscribed interests, which were explored in-depth using our BAP phenotyping protocol (described above) in members of unrelated large multiplex families primarily ascertained through the Collaborative Autism Study^33^. This in-depth characterisation was phenomenologically based^37^, whereby the number of traits within each domain was fully expanded through administration of the semi-structured interview (BAPI) with separate family members until no further traits were identified (saturation) to capture the entire range of BAP traits (Table 3).

**Table 3.** Identification of BAP traits through expansion of the BAP domains.

| **BAP domains** | **Expanded traits** |
| --- | --- |
| Communication<br>(speech and literacy) | Reduced capacity for clear narrative<br>Difficulty answering open ended questions<br>Reduced quantity of verbal output<br>Speech has little variation in tone (i.e. monotonous)<br>Unusual speech volume<br>Precise articulation and language<br>Use of an accent* |
| Social communication<br>(pragmatics and relationships) | An unusual or awkward greeting style<br>A limited capacity to develop rapport with assessors<br>Unusual eye gaze<br>Awkward social interactions<br>Making inappropriate or awkward comments either on history or during assessments<br>Tangential pragmatic style<br>Terse pragmatic style<br>Tendency to monologue rather than participate in reciprocal conversation<br>Opinionated in conversation<br>Overly technical language<br>Little appreciation of humour (during the Cartoon task)<br>Inflexible to intentional errors*<br>Tendency to anger easily<br>Narcissistic personality style<br>Self perception incongruent with views of others<br>Aloof personality style<br>Difficult or limited interpersonal relationships<br>Reduced affection<br>Reduced emotional empathy<br>Reduced cognitive empathy<br>Excessive worry |
| Circumscribed interests | Preference for structure in activities of daily living<br>Fastidious regarding personal appearance<br>Fastidious cleaning<br>Hobby or interest of unusual intensity, or restricted range of interests relative to peers<br>Large collections or hoarding of items<br>Focus on technicalities or minutiae<br>Recurrent thoughts (distressing)*<br>Recurrent thoughts (not distressing) |

\*BAP trait was removed from the final list (see text for details).

Initial phenotyping produced an exhaustive list of 36 BAP traits. Ordinal ratings of these traits were then assigned to capture subtle variations in their presentation, with severity rated on a scale of 0=absent, 1=mild, 2=moderate, and 3=severe. The presence of traits through each individual’s developmental history was also evaluated where available. Exploratory hierarchical cluster analysis was then performed to identify potential BAP endophenotypes. We used Ward’s method with Euclidean squared distances based on z-scores to progressively group traits by minimising the variability within clusters and maximising the variance between clusters^38^. Interpretation of cluster groupings was informed by the relative similarity and dissimilarity in the linkage output combined with clinical judgement, leading to the initial identification of five endophenotypes. Inspection of these endophenotypes revealed a consistent rating of 0 for two of the 36 traits across all interviews, leading to their removal. One further trait reflecting inflexibility to intentional errors was removed due to challenges reliably assessing it across interviewers, resulting in a final set of 33 BAP traits (Table 3).

#### Step 2: Validation of BAP Endophenotypes in Small Families

In the small families sample, an independent expert in ASD assessment (CG) interviewed and rated 45 participants on the 33 BAP traits based on all qualitative and quantitative data, with a subset (9%) rated via consensus between CG, IES and SJW to ensure consistency in ratings across both samples and to clarify borderline cases. As above, Ward’s hierarchical cluster analysis was used to examine natural trait groupings. This led to the identification of four endophenotypes that showed a high degree of similarity to the initial five cluster solution.

To account for a variable number of traits in each cluster we computed proportional scores, whereby scores on each trait (range 0-3) were summed and divided by the maximum total score for that cluster, to produce four cluster scores for each individual. An ROC curve was plotted for each cluster in the small families sample to identify optimum cut-off scores for determining endophenotypic status using Youden’s Index to allow mildly affected individuals to be included^39,40^. The highest score was used to represent the most prominent endophenotype for each individual, calculated as the difference between the observed endophenotype (i.e., cluster) score and the threshold score for the endophenotype (i.e., cut-off score).

#### Step 3: Assessment of BAP Endophenotypes in Family A and B

A team member who had not been involved in the phenotyping of Family A and B (Step 1) performed the endophenotype analysis (KT). Proportional scores for the four endophenotypes were calculated, and family members classified as having the endophenotype if their proportional score was greater than or equal to the cut-off scores identified in the small families analysis (step 2). As above, the highest score (observed endophenotype score – threshold endophenotype score) for any endophenotype was used to represent an individual’s most prominent endophenotype. A discriminant function analysis was then used to determine the sensitivity and specificity of the endophenotype approach to identifying the presence of the BAP in these families. In addition, endophenotype results were correlated with measures of intellect, executive, social and adaptive functions using conservative non-parametric Spearman’s correlations (*r*_*s*_).

## Results

### Hypothesis 1: Multiple individuals in large families demonstrate the BAP

Based on our rigorous protocol for phenotyping the BAP in large multiplex families, we identified 32 members with the BAP across Family A and B. Of the 23 members in Family A who did not have an ASD diagnosis, we detected the BAP in 17 (74%) individuals, with 6 individuals unaffected. In Family B, we detected 15 (63%) individuals with the BAP, with 9 individuals unaffected (Table 4).

**Table 4.**
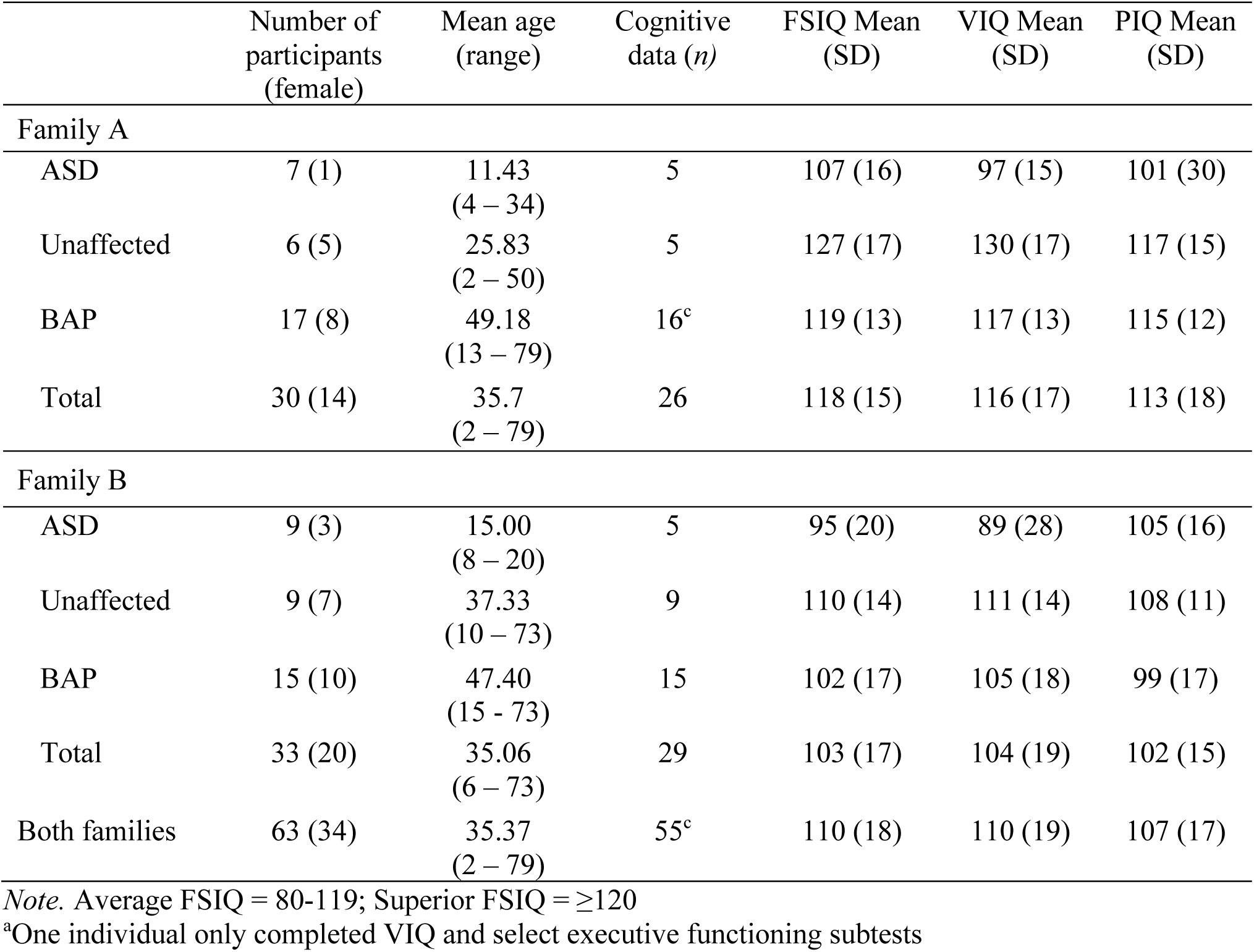
Intellectual functioning in Family A and B by diagnostic classification.

|  | Number of participants (female) | Mean age (range) | Cognitive data (*n*) | FSIQ Mean (SD) | VIQ Mean (SD) | PIQ Mean (SD) |
| --- | --- | --- | --- | --- | --- | --- |
| Family A |  |  |  |  |  |  |
| ASD | 7 (1) | 11.43<br>(4 – 34) | 5 | 107 (16) | 97 (15) | 101 (30) |
| Unaffected | 6 (5) | 25.83<br>(2 – 50) | 5 | 127 (17) | 130 (17) | 117 (15) |
| BAP | 17 (8) | 49.18<br>(13 – 79) | 16<sup>c</sup> | 119 (13) | 117 (13) | 115 (12) |
| Total | 30 (14) | 35.7<br>(2 – 79) | 26 | 118 (15) | 116 (17) | 113 (18) |
| Family B |  |  |  |  |  |  |
| ASD | 9 (3) | 15.00<br>(8 – 20) | 5 | 95 (20) | 89 (28) | 105 (16) |
| Unaffected | 9 (7) | 37.33<br>(10 – 73) | 9 | 110 (14) | 111 (14) | 108 (11) |
| BAP | 15 (10) | 47.40<br>(15 – 73) | 15 | 102 (17) | 105 (18) | 99 (17) |
| Total | 33 (20) | 35.06<br>(6 – 73) | 29 | 103 (17) | 104 (19) | 102 (15) |
| Both families | 63 (34) | 35.37<br>(2 – 79) | 55<sup>c</sup> | 110 (18) | 110 (19) | 107 (17) |

*Note.* Average FSIQ = 80-119; Superior FSIQ = $\geq 120$
<sup>a</sup>One individual only completed VIQ and select executive functioning subtests

Intellectual function was directly assessed in 54/63 (86%) individuals. Overall, participants were of average or greater intelligence. Average FSIQ was observed in 32/54 (59%) of individuals, while 20/54 (37%) demonstrated superior or very superior FSIQ (Table 4). We performed group-level comparisons of cognitive, social and adaptive functions between family members with and without the BAP using non-parametric and parametric tests (Mann-Whitney U and t-tests respectively), with the more conservative parametric tests reported here as there were no differences between these approaches. On average, individuals with the BAP demonstrated poorer pragmatic language, with significantly higher mean PRS scores (*M*=8.75, *SD*=7.08) compared to unaffected individuals (*M*=2.00, *SD*=3.05), *t*(40.99)=-4.42, *p*<0.001. No significant differences were observed for general intellect (FSIQ, VIQ, PIQ), the FPT, executive (D-KEFS) or adaptive function measures (ABAS-II, BRIEF).

### Hypothesis 2: Specific BAP endophenotypes exist across BAP domains

Based on our iterative characterisation process, four distinct endophenotypes of the BAP were reliably identified. Based on the natural grouping of traits, these reflected ‘socially unaware’, ‘pedantic’, ‘aloof’, and ‘obsessive’ endophenotypes (Table 5). At the highest level of the dendrogram of the 33 BAP traits there was a clear split, whereby traits of the socially unaware and pedantic endophenotypes were more similar to each other and more dissimilar to the combination of traits of the aloof and obsessive endophenotypes. There was a significant difference between the mean proportional scores of the unaffected and BAP groups, with the BAP group demonstrating significantly higher scores on all four endophenotypes (all *p*>0.015).

**Table 5.**
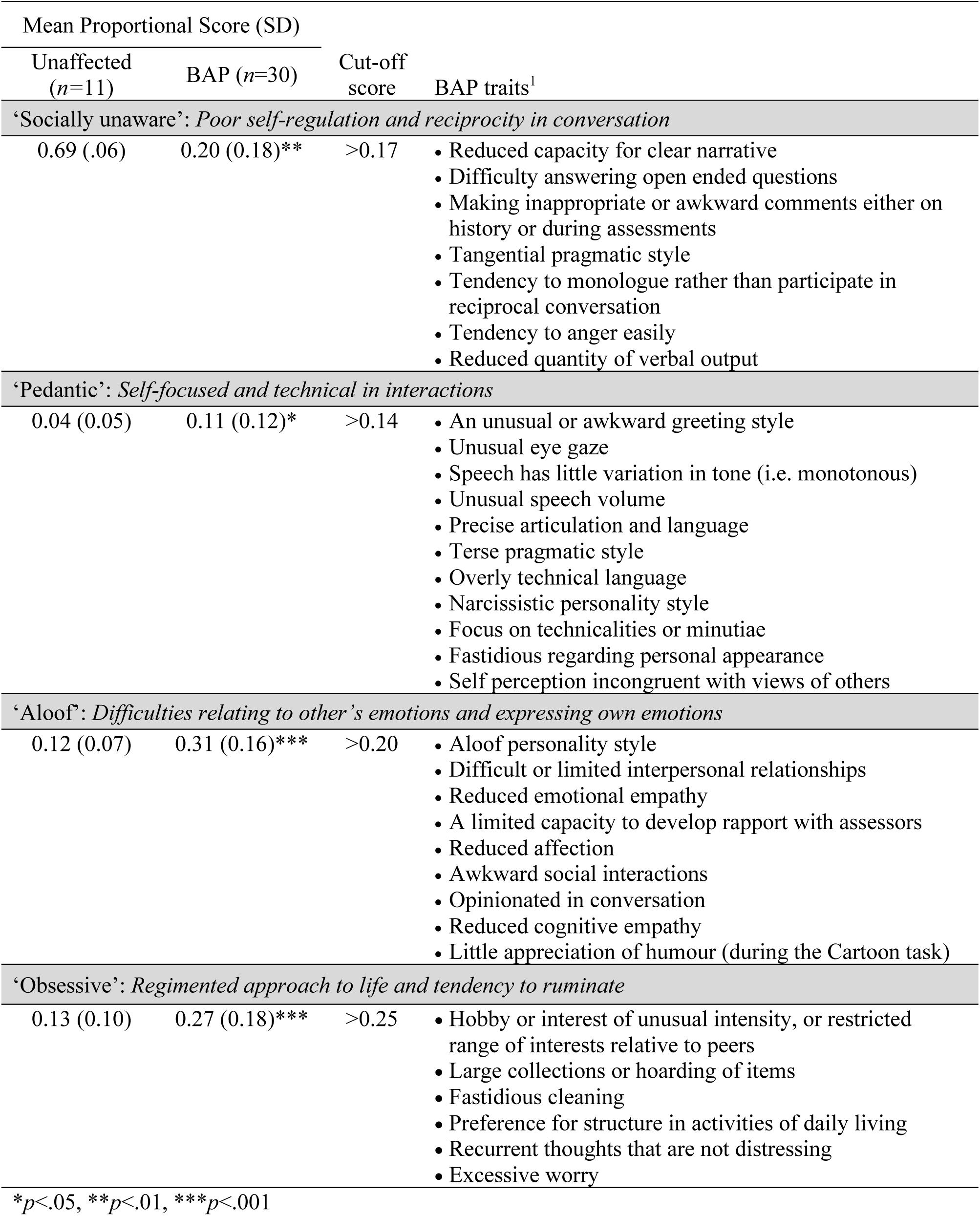
Four endophenotypes of the BAP in the small families sample.

Analysis of ROC curves indicated relatively good discrimination within the small families for the socially unaware, aloof and obsessive endophenotypes (all AUC > 0.73, all *p*<.025), and acceptable discrimination for the pedantic endophenotype (AUC=0.68, *p*=0.077). Although we note that Box’s M was violated in the discriminant function analysis (likely due to variation in the sample sizes), combined, the four endophenotypes captured 93% of cases (Wilk’s λ =0.47, χ^2^=27.83, *p*<.001). In particular, the endophenotypes showed high sensitivity for the BAP group (97%), characterised by higher proportional scores, and good specificity for the unaffected group (82%), with lower proportional scores (Table 5).

### Hypothesis 3: BAP endophenotypes vary in large multiplex families

Applying the above endophenotype thresholds to the proportional scores of the 33 BAP traits for members of Family A and B led to the identification of all individuals classified as having the BAP. Two additional BAP cases were identified in Family B based on the presence of above threshold endophenotype scores, indicating good utility of this approach (Fig.3). One individual was excluded from this analysis due to incomplete data (III-7). Across both families, the aloof endophenotype was most commonly observed (62%), followed by obsessive (60%), pedantic (55%) and socially unaware (48%). Approximately one quarter of family members met criteria for only one endophenotype, 15% met criteria for two, and the remainder met criteria for 3-4 (62%) (Fig.3). The dominant endophenotype across both families, as determined by the highest score, was aloof (47%), followed by obsessive (26%), socially unaware (18%) and pedantic (9%).

**Figure 3.**
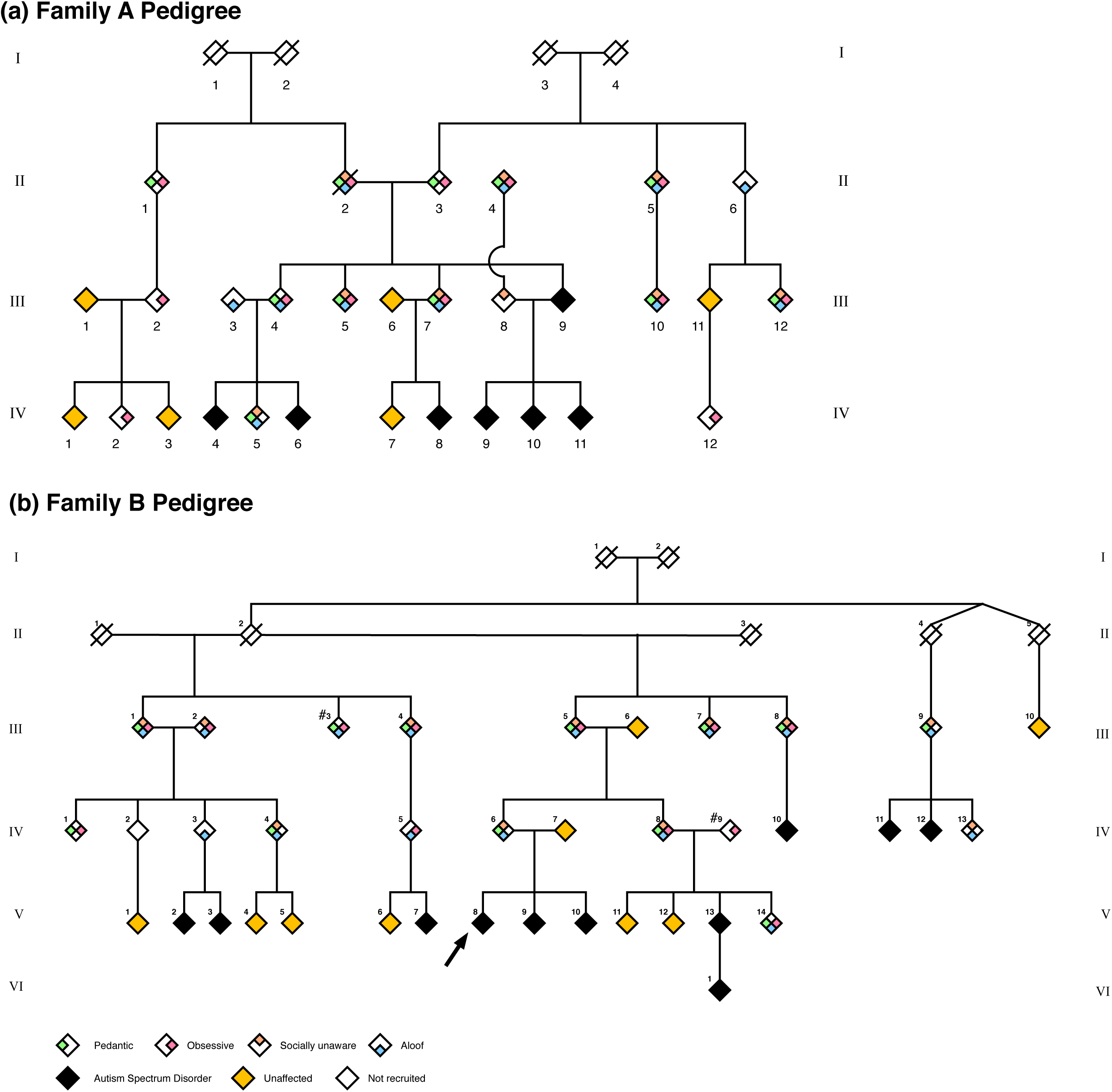
Scrambled pedigrees for Family A (panel A) and Family B (panel B) showing phenotypes and endophenotypes. All individuals with ≥ 1 endophenotype had the BAP, with the exception of two individuals from Family B (III-3 and IV-9) marked with an asterisk. These individuals were clinically determined as unaffected (Family B III-3 and IV-9) but had above threshold endophenotype scores based on ROC curves. Family members who were not phenotyped are not shown to preserve the anonymity of these families.

Family A appeared to have two endophenotype profiles, with one characterised by the presence of a single endophenotype (35%) seen in individuals who were mostly married-in (67%), contrasting with the second profile (41%) of all four endophenotypes, most evident in core family members (72%) (Fig.4). Overall, the obsessive endophenotype occurred most frequently (77%), followed equally by pedantic (65%) and aloof (65%), and then socially unaware (53%). The co-occurrence of the obsessive and pedantic endophenotypes was relatively common, seen in 29% of married-ins and core family members. Overall, there was a range of dominant endophenotypes across individuals, with aloof the most frequent (35%) particularly in core family members (83%).

**Figure 4.**
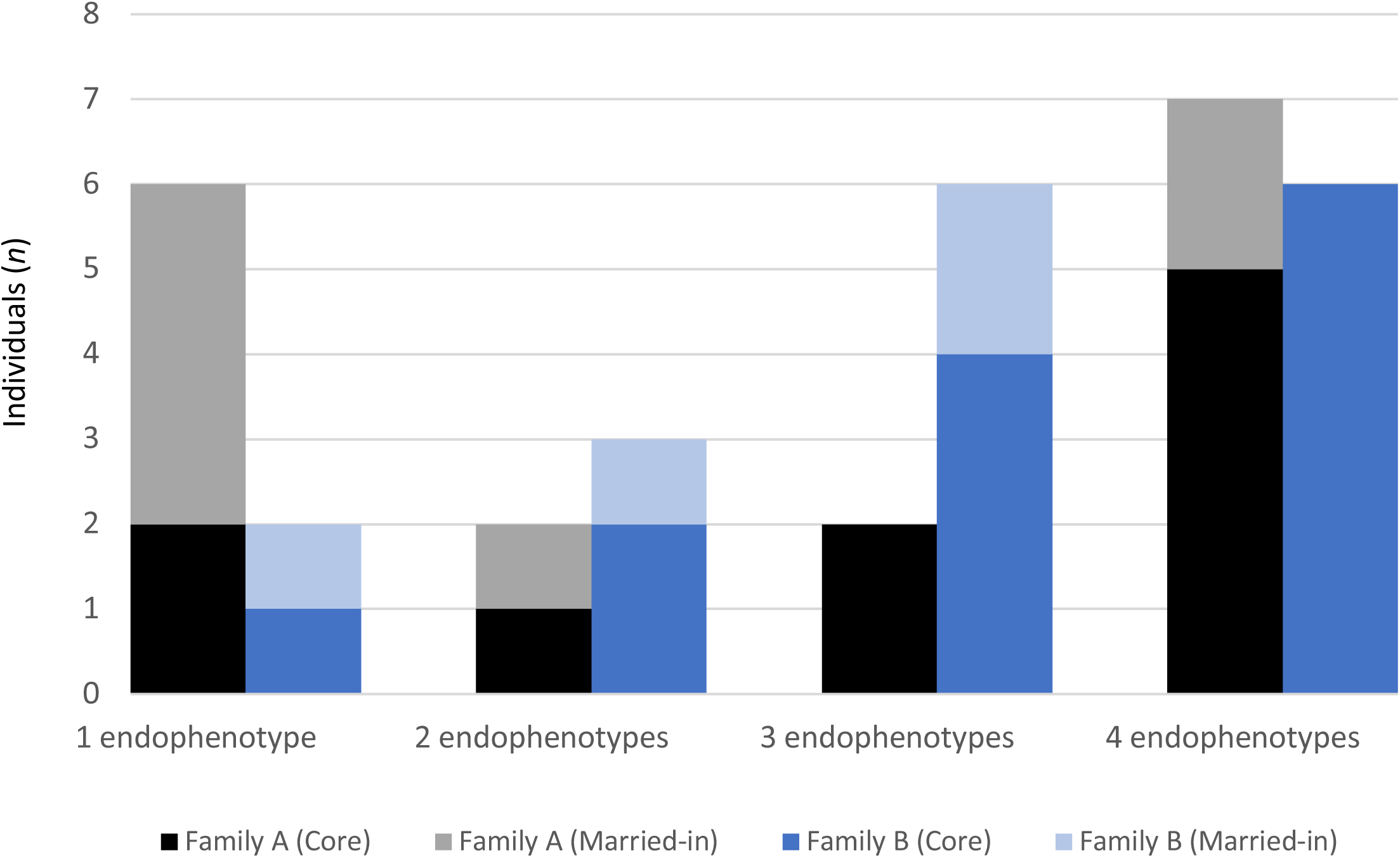
Number of BAP endophenotypes present in Family A and B. Individuals married-in to Family A tend to have a single endophenotype, indicating a more mild BAP presentation, in contrast with core family members who have multiple endophenotypes (obsessive most frequent). In Family B, married-in and core family members tend to have more than one endophenotype, with the aloof endophenotype most frequent.

Contrasting with Family A, Family B had more individuals (70%) with multiple endophenotypes, in both married-in and core family members (Fig.4). All four endophenotypes were again most frequently observed in core family members, indicative of a more severe BAP presentation. Unlike Family A, however, the aloof endophenotype occurred most frequently in Family B (88%), followed by obsessive (71%), pedantic (65%), and socially unaware (65%). The aloof endophenotype was also identified as dominant (59%), evident in 70% of core family members.

#### Correlates of the BAP Endophenotypes

Across both families, no sex or age differences were observed for any of the endophenotypes (all *p*<.200). Overall, a more severe BAP presentation (indicated by a greater number of endophenotypes) was associated with reduced social adaptive functioning on both self-report and objective measures of social communication (Table 6). In particular, a more severe BAP presentation showed a strong correlation with more severe pragmatic language difficulties, with scores for each endophenotype also significantly correlated. A similar relationship was evident for the ability to detect a faux pas in social discourse and self-reported social functioning, particularly for family members with the socially unaware endophenotype (Table 6).

**Table 6.**
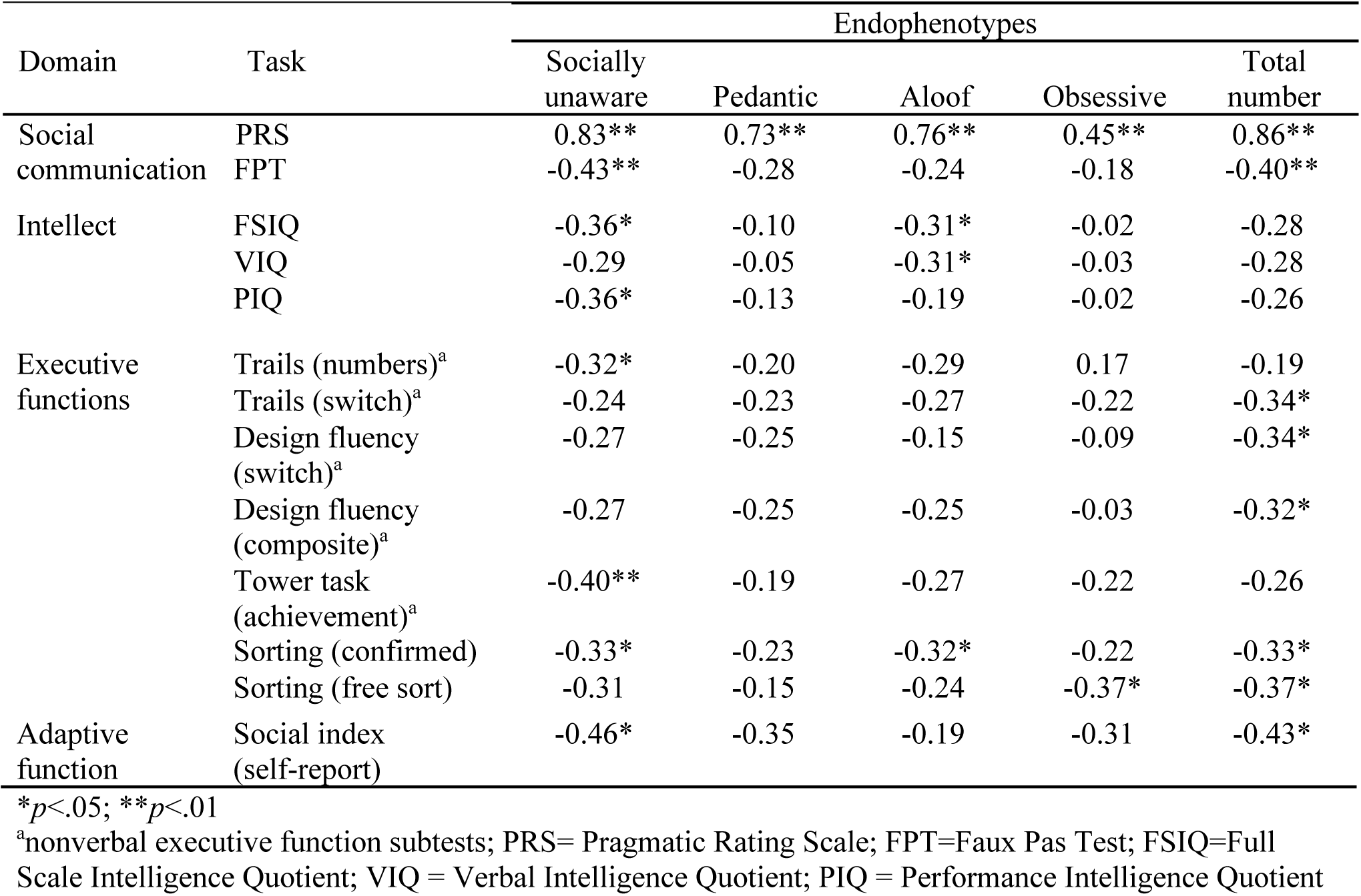
Correlations between endophenotypes and quantitative measures in Family A and B.

For the cognitive measures, a more severe BAP presentation was associated with reduced executive functioning, particularly for nonverbal measures of cognitive flexibility (switching and fluency; Table 6). A pattern of weaker correlations was also evident for specific endophenotypes, including lower IQ in the socially unaware and aloof endophenotypes (Table 6).

## Discussion

We studied the BAP to highlight the phenotypic variation within and between high-risk ASD families, to improve identification of individuals crucial for accurate molecular genetic analysis. We identified multiple individuals with the BAP in large multiplex families using rigorous phenotyping and a new endophenotyping approach, which was validated in an independent sample of small ASD families. The results of this work show that specific BAP endophenotypes exist across the traditional BAP domains of social relationships, communication, and circumscribed interests and behaviour, providing a more nuanced way to detect subtle features of the BAP. Moreover, these endophenotypes show different patterns of inheritance in two large multiplex families, supporting their use to identify autism genes of dominant effect.

Despite major advances in ASD genetics, aetiology in the majority of cases remains unknown. The research model employed here to phenotype rare large multiplex families reveals a pattern consistent with autosomal dominant inheritance of ASD/BAP traits that would not have been captured without such rigorous phenotyping. Fifteen individuals (23%) met criteria for ASD and 33 (51%) the BAP, including some married-in individuals. Our promising endophenotype analysis provides further insight into specific profiles of the BAP and its varied presentation. Traditionally, ASD family studies include 2-3 affected individuals^41,42^. For example, four candidate ASD genes were identified in seven ASD/BAP pedigrees with ≥3 affected individuals^43^. Larger multiplex families remain scarce in the literature^20,44^. Here, we identified more subtle indicators of carrier status in two large families, using a robust endophenotyping method with good sensitivity and specificity to detect the BAP in two independent samples.

### Endophenotypes of the BAP

Over the last 20 years, the BAP has emerged as strongly associated with ASD. The BAP is considered a marker of carrier status of genes that may contribute to autism risk^21,45^. Here we aimed to dissect the BAP into endophenotypes to understand the phenotypic variation within and between families. Importantly, each endophenotype cluster was characterised by a combination of communication, personality and behavioural indicators showing how specific traits across the traditional BAP domains may group together to form distinct endophenotypes or ‘profiles’. As summarised in Table 7, these profiles capture identifiable ‘personas’ that have core characteristics with high face validity. These profiles also vary with functional correlates in distinct ways, supporting their construct validity. For example, the aloof endophenotype was characterised by a lack of innate social motivation or ability to meaningfully connect and empathise with others, associated with decreased theory of mind, lower executive and intellectual functioning. One individual dominant for the aloof endophenotype described social interactions as “a means to an end”. In contrast, the pedantic endophenotype was primarily characterised by detail-oriented traits, showing no associations with intellectual, executive or adaptive functions. Unsurprisingly, given the importance of social communication deficits in ASD and the BAP, all endophenotypes were associated with poor social communication, with the socially unaware endophenotype most broadly affected across social, intellectual, executive and adaptive function domains (Table 7).

**Table 7.**
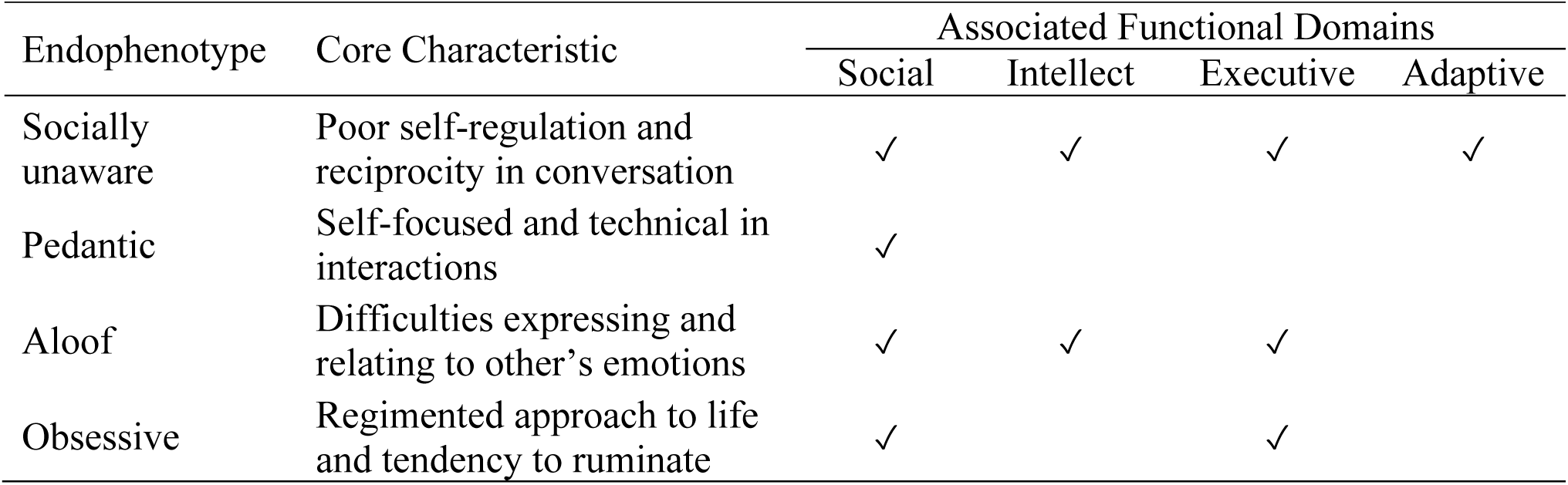
Summary of the BAP endophenotypes and their functional correlates.

By clustering traits across traditional BAP domains, endophenotype profiles may improve detection of the BAP and thus, advance gene discovery. Specifically, in contrast with a traditional domain approach, where an individual may show mild BAP features across all domains but fail to meet criteria, an endophenotype approach allows individuals with autism susceptibility genes to be captured by meeting threshold criteria for a specific profile. Importantly, replication and validation of the proposed BAP endophenotypes is needed, using targeted assessments to further validate and refine the traits characterising each endophenotype. This, in turn, will provide the foundation for more efficient assessment protocols, and more sophisticated and granular mapping of psychological and neural correlates where results have been mixed to date^46^.

### Careful endophenotyping will enable genetic insights

Consistent with previous literature, phenotypic heterogeneity was evident in both families at the endophenotype level suggesting a single familial mutation may produce a phenotypic spectrum, with other genetic, epigenetic and environmental factors influencing expression. With the advancement of high-throughput next generation sequencing technologies, meticulous phenotypic characterisation of both affected and apparently unaffected individuals remains essential for accurate data interpretation. In other words, identification of subtle endophenotypes, such as the four identified here, are crucial for advancing gene discovery programs.

Although multiplex families with ASD are genetically homogeneous, our phenotyping analysis suggests possible bi-lineal inheritance of the BAP in both families. Therefore, multiple risk alleles may contribute to ASD/BAP in later generations, consistent with recent genetic and phenotyping evidence^47,48^. The importance of unique *de novo* genetic changes in both sporadic (or ‘simplex’), ASD^17^, and small multiplex ASD families^44^ has become increasingly apparent. However, with at least seven individuals with ASD and many more with the BAP in our families, there is less likelihood of *de novo* changes contributing to each phenotype. It is much more likely that there is a single genetic variant of major phenotypic effect in each family, with the possibility that there are additional *de novo* genetic changes in some individuals that contribute to phenotypic severity.

### Limitations

The intensive nature of the study meant that clinicians were not blinded to family relationships, potentially leading to investigator bias. However, our diagnostic method of consensus between experienced clinicians aligns with current best practice for ASD/BAP diagnosis and was informed by quantitative and qualitative measures. We selected a relatively low threshold for BAP classification, leading to the identification of many affected individuals. However, this approach is justified in a family with a clear genetic liability for ASD and was validated by the finding of consistent data-driven endophenotypes in the small families. Successful gene identification in future work requires capture of all individuals who may carry the putative variant, with the approach outlined here enabling more robust gene identification work.

## Conclusion

Despite significant advances in unravelling the heterogeneity of ASD, in most cases, the underlying genetic aetiology remains unknown in part due to difficulties identifying endophenotypes and potential carriers. We used a rigorous phenotyping approach to characterise the BAP in two large multiplex families with dominant inheritance of ASD and the BAP. Further phenotypic delineation identified four endophenotypes, showing differentiation of BAP features beyond traditional domain approaches. This endophenotype approach advances our understanding of the phenotypic spectrum to improve detection of the BAP in research and clinical practice, facilitating gene discovery, neuroimaging investigations, and psychological studies.

